# Counting the paternal founders of Austroasiatic speakers associated with the language dispersal in South Asia

**DOI:** 10.1101/843672

**Authors:** Prajjval Pratap Singh, Shani Vishwakarma, Gazi Nurun Nahar Sultana, Arno Pilvar, Monika Karmin, Siiri Rootsi, Richard Villems, Mait Metspalu, Doron M Behar, Toomas Kivisild, George van Driem, Gyaneshwer Chaubey

**Affiliations:** Cytogenetics Laboratory, Department of Zoology, Banaras Hindu University, Varanasi 221005, India; Estonian Biocentre, Institute of Genomics, University of Tartu, Tartu 51010, Estonia; Centre for Advanced Research in Sciences (CARS), DNA Sequencing Research Laboratory, University of Dhaka, Dhaka, 1000, Bangladesh; Statistics and Bioinformatics Group, School of Fundamental Sciences, Massey University, Palmerston North 4410, New Zealand; Department of Human Genetics, Katholieke Universiteit Leuven, Leuven, 3000, Belgium; Institut für Sprachwissenschaft, Universität Bern, 3012 Bern, Switzerland; Sydney Social Sciences and Humanities Advanced Research Centre, University of Sydney, Australia

**Keywords:** Austroasiatic, South Asia, Haplogroup O2, Mundari, Tibeto-Burman

## Abstract

The phylogenetic analysis of Y chromosomal haplogroup O2a-M95 was crucial to determine the nested structure of South Asian branches within the larger tree, predominantly present in East and Southeast Asia. However, it had previously been unclear how many founders brought the haplogroup O2a-M95 to South Asia. On the basis of the updated Y chromosomal tree for haplogroup O2a-M95, we analysed 1,437 male samples from South Asia for various downstream markers, carefully selected from the extant phylogenetic tree. With this increased resolution, we were able to identify at least three founders downstream to haplogroup O2a-M95 who are likely to have been associated with the dispersal of Austroasiatic languages to South Asia. The fourth founder was exclusively present amongst Tibeto-Burman speakers of Manipur and Bangladesh. In sum, our new results suggest the arrival of Austroasiatic languages in South Asia during last five thousand years.

## Introduction

Among the four major language families present in South Asia, the Austroasiatic language family has the smallest number of speakers. Nevertheless, its geographical location and exclusive distribution amongst tribal populations make Austroasiatic one of the most intriguing language families in the context of the Subcontinent ^1–3^ where it is represented by the major Mundari and minor Khasi branches. ^4^ The advancement of technology has helped to resolve the long standing debate about the arrival of Austroasiatic (Mundari) speakers to South Asia. The three types of genetic markers (mtDNA, Y chromosome and autosomes) revealed the extraordinary migratory phenomenon. Because of the incompatible histories of their paternal and maternal ancestries, the origin and dispersal of Austroasiatic language in these populations had remained a debatable issue for many years.^5,6^ For Mundari speakers, the exclusive South Asian maternal ancestry pinpointed their origin in South Asia, whereas their overwhelming East/Southeast-Asian-specific paternal ancestry associated with haplogroup O2a-M95 suggested a dispersal from East to West. ^3,6–8^ Another Austroasiatic branch, Khasi, evinced mixed ancestries for both the paternal and maternal lines. ^9^ Over a century of ethnographical and Indological scholarship had identified Austroasiatic language communities as representing the earliest settlers of South Asia, ^3,7,9–12^ so that this view represented the consensus. ^13^ Then, the increasing resolution of the Y-chromosomal tree and Y-STR-based coalescent calculations began to challenge our view of these populations. Despite the problems with STR-based coalescent calculations, the Y-STR variance for haplogroup O2a-M95 was consistently reported to be significantly higher in Southeast Asia. ^8,14^

With the technological advancement, a much clearer picture has now emerged, showing a strong association of largely male mediated gene flow for the dispersal of Mundari speakers from Southeast Asia to South Asia. ^14–16^ The autosomal data showed a bidirectional geneflow between South and Southeast Asia. ^14^ In addition, the global distribution and internal phylogeny of Y-chromosomal haplogroup O2a-M95 decisively demonstrated that this language dispersal transpired from Southeast Asia to South Asia. In a recent genome-wide study of a large number of South and Southeast Asian populations, Dravidian speakers of Kerala and Lao speakers from Laos were found to be the best ancestral proxies for Mundari speakers. ^15^ The admixture dates for these ancestries were estimated to be between 2-3.8 kya. Moreover, this study also found clear-cut genetic differentiation between North and South Munda groups ^15^.

Based on the prevalence of haplogroup O2a-M95, the arrival of Mundari speakers to South Asia was shown to have been largely male-mediated. ^6,8,14^ However, it remained unclear how many paternal founders had brought this haplogroup to South Asia. The recent admixture time estimated through autosomal data has now narrowed down the arrival of Mundari ancestors to South Asia. ^15^ Therefore, a marker such as M95, which originated >12 kya, would not help us to understand the most recent founding lineages. The association of haplogroup O2a-M95 with the arrival of Mundari speakers has been explored previously in detail. ^6,8,10,14^ However, the lack of downstream markers made it impossible to discern the number of most recent founders who had originally borne the language to South Asia. Moreover, previous studies on Tibeto-Burman populations of Northeast India and Bangladesh also reported this same haplogroup,^6,8,10,14,17^ giving rise to the contentious question as to whether the expansion of Tibeto-Burmans could have involved the assimilation of Austroasiatic populations. Therefore, we took advantage of recent technological advancements and extracted the downstream SNPs associated with Munda-related populations from one of our recent studies. ^18^ We genotyped these SNPs for large number of Mundari, Tibeto-Burman and Bangladeshi population groups (Fig. 1) and reconstructed the phylogeographical distribution of downstream branches of haplogroup O2a-M95 in South Asia.

**Figure 1.**
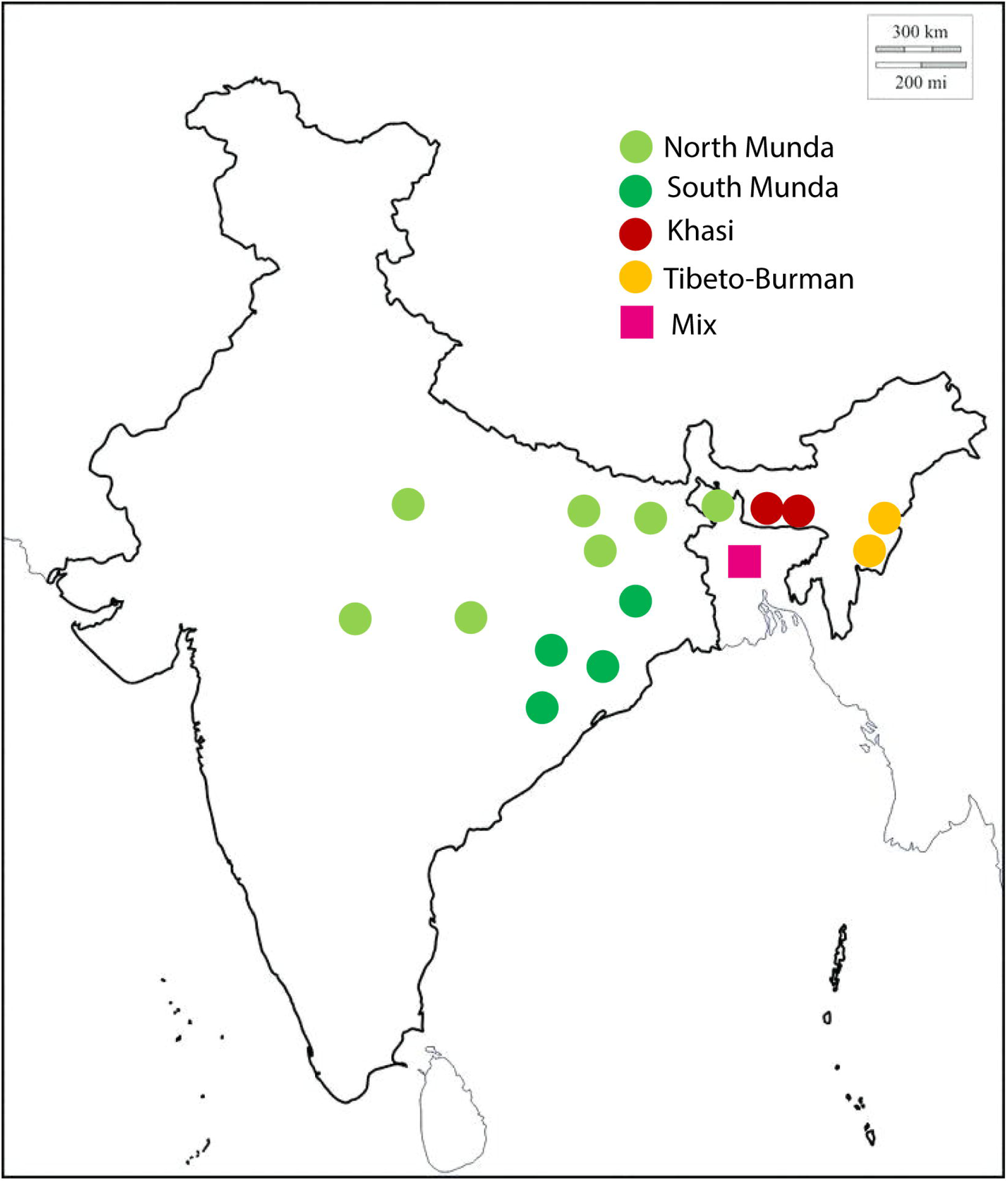
The sampling locations of various ethnic groups used in the present analysis. The detailed linguistic information of all the Bangladeshi samples genotyped were not present, therefore we have designated them as ‘mix’ group.

## Methods

In our quest to identify the founder branches of haplogroup O2a-M95 who are likely to have introduced Austroasiatic languages to South Asia, we first extracted the Y chromosome sequences from haplogroup O2a-M95 derived samples ^18,19^. We note that from South Asia, the 1000 genome samples and Karmin et al. ^18^ both had two South Asian individuals, however, both of the 1000 genome samples were falling in a single branch ^19^, whereas, Karmin et al. samples were present on two distinct branches ^18^. Therefore, we used Karmin et al. data and reconstructed phylogenetic trees of haplogroup O2a-M95 based on the ~10MB region of the Y chromosome. ^18^ We extracted SNPs focussing on those branches where at least one South Asian sample was present. Thereafter, we designed primers targeting selected SNPs and genotyped large number of samples either via RFLP (restriction fragment length polymorphism) or Sanger sequencing (Supplementary Table 1). We first genotyped the M95 marker among all the samples (n=1437) and followed the hierarchal genotyping for the remaining SNPs among the M95 derived samples. We used the data generated in the present study and reconstructed the phylogeny of haplogroup O2a-M95 with the frequency of each and every SNP among selected populations (Fig. 2).

**Figure 2.**
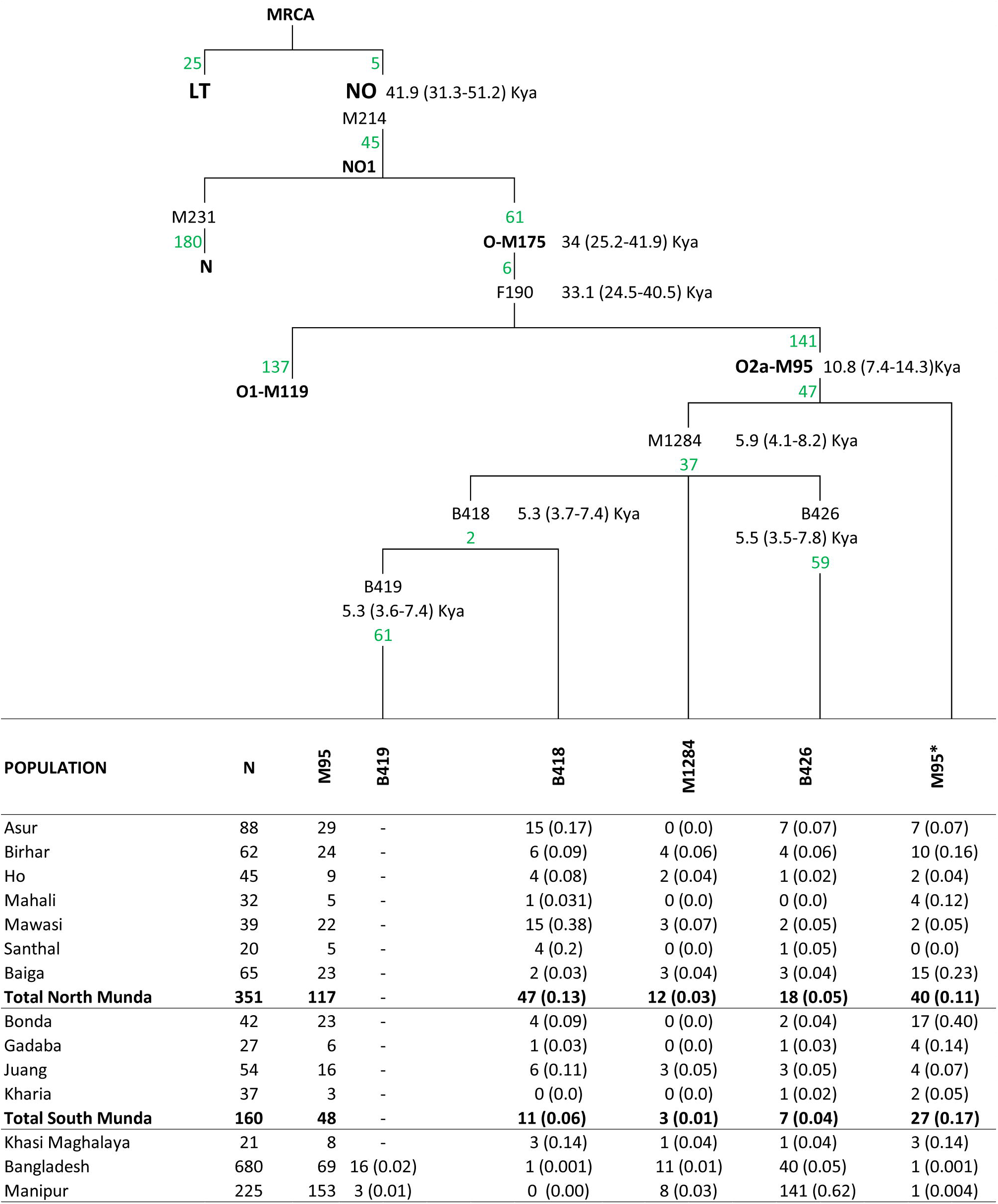
The updated phylogeography of haplogroup O2a-M95 in South Asia. The branch coalescent times were calculated according to Karmin et al. ^18^. The number of mutations of a branch are shown in green colour. For each branch, the total number of individuals of a population as well as the frequency are shown. The M95* is M95x(B418, B419, B426, M1284).

For the coalescent age estimation of various nodes, we have used the mutation rate calibrations, 0.73×10^−9^/bp/year (95% CI 0.63–0.95 ×10^−9^/bp/year) published elsewhere ^18^. By the inclusion of six new markers downstream of M95, we were able to resolve the incompatible dates of the autosomal and Y chromosomal studies ^14,15^ as well as obtain a clearer picture of the phylogeography of haplogroup O2a-M95 specific to South Asia.

## Results and Discussion

The newly reconstructed South-Asian-specific phylogeny of haplogroup O2a-M95 is now defined by four new more recent branches showing a coalescent time in the range of 5 kya (kilo years ago) (Fig. 1). These four founders gave rise to ~77% of individuals carrying haplogroup O2a-M95 in South Asia, and the distribution of these four founders amongst the groups studied is intriguing. Founder one (represented by marker B419) is virtually absent amongst Mundari speakers and is instead exclusively found amongst Tibeto-Burman speaking population in Manipur and Bangladesh (Figs. 2,3). Conversely, Founder two (marker B418) occurs most frequently in North Munda language communities and is absent among Manipuri populations. Founder three (marker M1284) is infrequent, yet nevertheless present amongst all the groups studied here (Fig. 3). Founder four (marker B426) is also present amongst all the groups and, most importantly, appears to be one of the leading founders of haplogroup O2a-M95 amongst those Tibeto-Burman populations where this paternal lineage is found (Figs. 2,3). With these enhancements, we were able to assign 59% of Mundaris, 62% of the Khasis, and 99% of Manipuri and Bangladeshi individuals to the four downstream branches of haplogroup M95-O2a, which have originated in the course of last five thousand years (Fig. 1). We designated unclassified O2a-M95 derived samples as the M95* which is phylogenetically M95x(B418, B419, B426, M1284).

**Figure 3.**
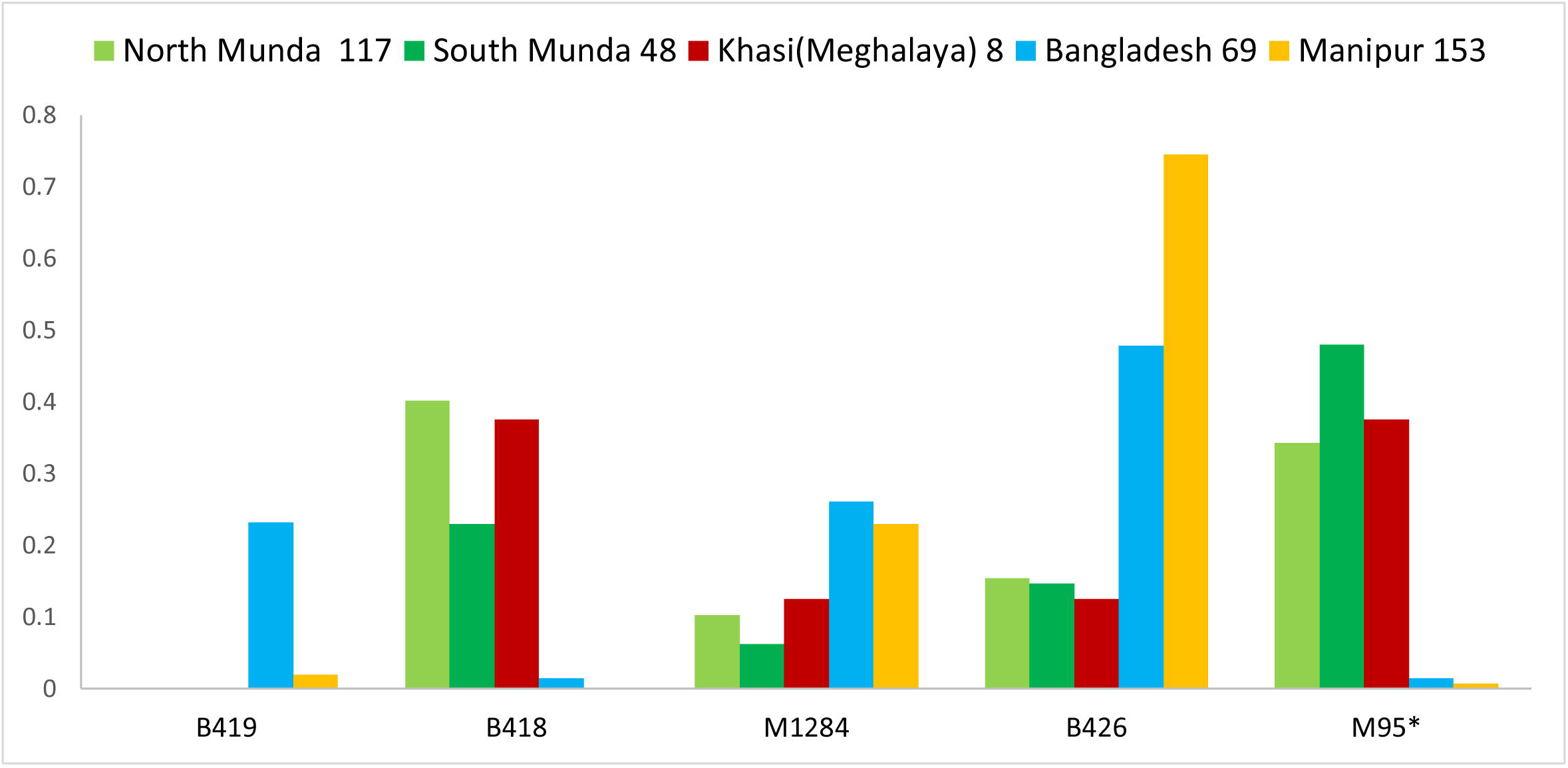
The bar plot showing the frequency of M95* and the associated branches amongst various ethnic groups of South Asia. This frequency is calculated out of the extant samples falling in haplogroup O2a-M95.

Our current phylogenetic understanding suggests that haplogroup O2a-M95 originated >30 kya in East/Southeast Asia and expanded more recently into South Asia (Fig. 2). The frequency distribution and coalescent pattern of various founders amongst Mundari groups clearly show that the migration of Austroasiatics to South Asia was much later than the origin of O2a-M95 (Fig. 2). The diverse founders as well as the large number of unclassified samples (41% for Mundari, 38% for Khasi and 1% for Tibeto-Burmans) suggest that the migration of Austroasiatic speakers to South Asia was not associated with the migration of a single clan or a drifted population. Neither does the contrasting distribution of various founders discovered in this study amongst both Mundari and Tibeto-Burman populations support the assimilation of the former to the latter.

Thus, with the help of improved resolution, we have upgraded the phylogeography of haplogroup O2a-M95 and identified at least four founders in South Asia. Among the four founders identified by this study, one was exclusively present amongst Tibeto-Burman populations of Manipur and Bangladesh. The sequencing of a few representative samples from unresolved M95 samples would, it is hoped, be able to resolve this matter more fully in the near future.

## Ethical approval

This study was approved by the regional ethical committee of Institute of Science, Banaras Hindu University, India (I.Sc./ECM-XII/2018-19/06).

## Acknowledgements

This work is supported by the National Geographic Explorer grant HJ3-182R-18. PPS is supported by the CSIR-JRF doctoral fellowship. MK, SR, RV, MM, DMB and TK are supported by institutional research funding IUT (IUT24-1) of the Estonian Ministry of Education and Research. MK is supported by the European Union through the European Regional Development Fund (Project No. 2014-2020.4.01.15-0012) The funders had no role in study design, data collection and analysis, the decision to publish or the preparation of the manuscript.

## Compliance with ethical standards Conflict of interest

The authors declare that they have no conflict of interest.

## References

1. Bellwood, P. The Austronesian dispersal and the origin of languages. Sci Am 265, 88–93 (1991).

2. Diffloth, G. The contribution of linguistic palaeontology to the homeland of Austro-Asiatic. in The peopling of East Asia (eds. Sagart, L., Blench, R. & Sanchez-Mazas, A.) 323 (Routledge, 2005).

3. Kumar, V. & Reddy, M. Status of Austro-Asiatic groups in the peopling of India: An exploratory study based on the available prehistoric, linguistic and biological evidences. J Biosci 28, 507–522 (2003).

4. Driem, G. van. Languages of the Himalayas. (2001).

5. Chaubey, G., Metspalu, M., Kivisild, T. & Villems, R. Peopling of South Asia: investigating the caste-tribe continuum in India. BioEssays News Rev. Mol. Cell. Dev. Biol. 29, 91–100 (2007).

6. Sahoo, S. et al. A prehistory of Indian Y chromosomes: evaluating demic diffusion scenarios. Proc Natl Acad Sci U A 103, 843–8 (2006).

7. Basu, A. et al. Ethnic India: a genomic view, with special reference to peopling and structure. Genome Res 13, (2003).

8. Sengupta, S. et al. Polarity and temporality of high-resolution y-chromosome distributions in India identify both indigenous and exogenous expansions and reveal minor genetic influence of Central Asian pastoralists. Am. J. Hum. Genet. 78, 202–21 (2006).

9. Reddy, B. M. et al. Austro-Asiatic tribes of Northeast India provide hitherto missing genetic link between South and Southeast Asia. PLoS ONE 2, e1141 (2007).

10. Kumar, V. et al. Y-chromosome evidence suggests a common paternal heritage of Austro-Asiatic populations. BMC Evol Biol 7, 47 (2007).

11. Majumder, P. P. & Basu, A. A genomic view of the peopling and population structure of India. Cold Spring Harb. Perspect. Biol. 7, a008540 (2014).

12. Malhotra, K. Morphological composition of the people of India. J. Hum. Evol. 45–53 (1978).

13. Chakravarti, A. Human genetics: Tracing India’s invisible threads. Nature 461, 487–8 (2009).

14. Chaubey, G. et al. Population Genetic Structure in Indian Austroasiatic speakers: The Role of Landscape Barriers and Sex-specific Admixture. Mol. Biol. Evol. 28, 1013–24 (2011).

15. Tätte, K. et al. The genetic legacy of continental scale admixture in Indian Austroasiatic speakers. Sci. Rep. 9, 3818 (2019).

16. Metspalu, M., Mondal, M. & Chaubey, G. The genetic makings of South Asia. Genet. Hum. Orig. 53, 128–133 (2018).

17. Trivedi, R. et al. Genetic imprints of pleistocene origin of indian populations: a comprehensive Phylogeographic sketch of Indian Y-chromosomes. Int J Hum Genet 8, (2008).

18. Karmin, M. et al. A recent bottleneck of Y chromosome diversity coincides with a global change in culture. Genome Res. (2015).

19. Poznik, G. D. et al. Punctuated bursts in human male demography inferred from 1,244 worldwide Y-chromosome sequences. Nat. Genet. (2016).

